# Multi-photon attenuation-compensated light-sheet fluorescence microscopy

**DOI:** 10.1101/2019.12.12.873877

**Authors:** Madhu Veettikazhy, Jonathan Nylk, Federico Gasparoli, Adrià Escobet-Montalbán, Anders Kragh Hansen, Dominik Marti, Peter Eskil Andersen, Kishan Dholakia

## Abstract

Attenuation of optical fields owing to scattering and absorption limits the penetration depth for imaging. Whilst aberration correction may be used, this is difficult to implement over a large field-of-view in heterogeneous tissue. Attenuation-compensation allows tailoring of the maximum lobe of a propagation-invariant light field and promises an increase in depth penetration for imaging. Here we show this promising approach may be implemented in multi-photon (two-photon) light-sheet fluorescence microscopy and, furthermore, be achieved in a facile manner utilizing a graded neutral density filter, circumventing the need for complex beam shaping apparatus. A “gold standard” system utilizing a spatial light modulator for beam shaping is used to benchmark our implementation. The approach will open up enhanced depth penetration in light-sheet imaging to a wide range of end users.

## 1. INTRODUCTION

Light-sheet fluorescence microscopy (LSFM) has transformed the field of imaging in recent years, owing to its optical sectioning capabilities, resulting in fast, highly resolved images with significantly reduced photo-bleaching and photo-toxicity [1, 2]. Propagation-invariant light fields, such as Airy and Bessel beams, have been employed in LSFM not only because of their pseudo-nondiffracting properties which enables them to retain their transverse profile on propagation, but also due to their self-healing capabilities on interaction with obstacles during propagation [3–6]. However, attenuation due to scattering and absorption results in an exponential decay of intensity of any given optical field as it penetrates deep into tissue and limits the penetration depth achievable for deep tissue imaging. Recently, the capability to shape the envelope profile of a light field arbitrarily [7–10] has been demonstrated to counteract the attenuation-induced exponential decrease in intensity, by tailoring an exponential rise in intensity along the direction of propagation [9]. Building upon this, LSFM exploiting attenuation-compensated Airy beams in single-photon imaging has demonstrated improved image quality at depth in attenuating biological specimens without any increase in the peak intensity of the illuminating light-sheet [11]. This is achieved by the selective delivery of additional intensity to greater depths within the attenuating medium, potentially minimizing photo-damage across the specimen. Our previous work utilized a spatial light modulator (SLM) for generation of attenuation-compensated Airy beams solely for single-photon imaging. While this approach offers excellent beam quality and has the flexibility to dynamically adjust the beam shape to optimally counteract the specimen attenuation, it adds cost and complexity to such a system, limiting the potential uptake of the method.

In this work, we show that attenuation-compensation of propagation invariant Airy fields can be achieved for multi-photon (two-photon) LSFM, and can be implemented in an inexpensive and facile manner. This is achieved by exploiting readily-available graded neutral density filters (NDF), effectively eliminating the need for an SLM. An increase in SNR of up to 45% is observed with NDF-based two-photon attenuation-compensated Airy LSM in biological specimens. Additionally, we demonstrate this NDF-based approach in single-photon LSFM.

## 2. MATERIALS AND METHODS

The cylindrical pupil function of an Airy light-sheet, compensated to overcome exponential intensity decay is represented as [11]

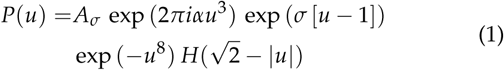

where *u* is the normalized pupil coordinate corresponding to the *z*–axis of the microscope (see Fig. 1(a)), *A*_*σ*_ is a real scaling factor, *α* controls the propagation-invariance of the Airy light-sheet [5], *σ* dictates the degree of linear attenuation-compensation, *H*(.) is the Heaviside step function, and the light-sheet propagates in the positive *x* direction. The top line of Eq. (1) describes the pupil function required for attenuation-compensation, the bottom line describes the transverse envelope of the beam and the use of different envelope functions will still yield attenuation-compensated Airy beams. The only modification required to transform an Airy beam into an attenuation-compensated Airy beam is the addition of an exponential amplitude term (exp(*σ* [*u* − 1])).

**Fig. 1.**
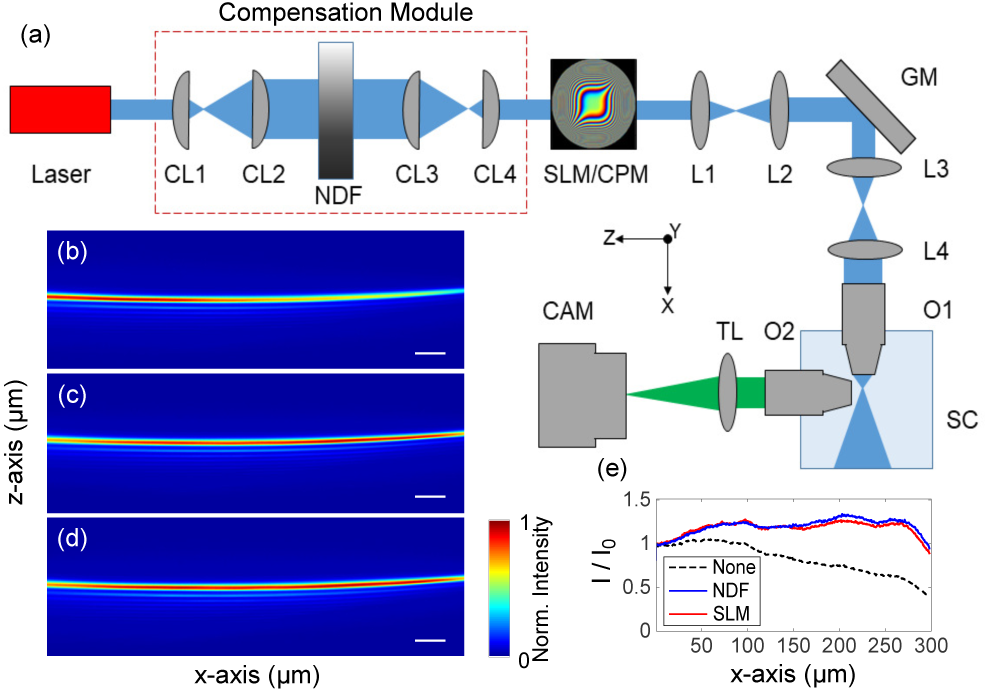
Experimental design of the attenuation-compensated Airy light sheet microscope, and Airy beam profiles. (**a**) Experiment setup with attenuation-compensation provided by NDF. (b - d) show the 2+1D Airy light-sheet profiles attenuated by an absorbing NIR dye (2 mM) and visualized in fluorescein to show two-photon fluorescence with (**b**) no attenuation-compensation, (**c**) attenuation-compensated by NDF, and (**d**) attenuation-compensated by SLM (*σ* = 0.5). (**e**) Normalized peak transverse intensity versus propagation distance corresponding to beam profiles in (b - d). CAM: Camera, CL: Cylindrical Lens, CPM: Cubic Phase Mask, GM: Galvo Mirror, L: Lens, NDF: Neutral Density Filter, O: Objective, SC: Sample Chamber, SLM: Spatial Light Modulator, TL: Tube Lens. Scalebars: (b - d) 20 µm.

Our two-photon attenuation-compensated Airy LSFM is based on the OpenSPIM design [12] in a digitally scanned light-sheet microscope (DSLM) configuration [13] as shown in Fig. 1(a). The Ti:Sapphire ultrashort pulsed laser (Coherent Chameleon Ultra II, central wavelength 810 nm, 140 fs pulse duration, 80 MHz repetition rate) is spatially filtered and expanded before being directed onto a spatial light modulator (SLM; Hamamatsu, LCOS X10468-04) or cubic phase mask (CPM; PowerPhotonic, custom LightForge mask) to generate the phase profile required to generate a 2+1D Airy profile [11, 14]. The SLM (CPM) is then imaged onto a galvo mirror (GM; Thorlabs GVS001) by lenses L1 and L2 and then onto the back aperture of the illumination objective (O1; Nikon, N10XW-PF, 0.30/10x, water immersion) by lenses L3 and L4.

Attenuation-compensation can be implemented on the SLM or by the addition of the compensation module (red dashed box in Fig. 1(a)). The compensation module comprises a linear graded neutral density filter (NDF; Thorlabs, NDL-25C-4, Optical density: 0.04 - 4.0) and two cylindrical telescopes oriented to elongate the beam along the *z*–axis (CL1 and CL2) incident onto the NDF and then contract it to the original beam dimensions (CL3 and CL4). The focal lengths of the cylindrical lenses were chosen to give magnifications of 4x and 0.25x respectively. The compensation module is positioned such that there is an imaging relation between the NDF and the SLM/CPM along the *z*–axis. Different sizes of beam incident on the NDF will yield different intensity (amplitude) gradients across the beam and therefore achieve different strengths of intensity modulation along the beam. For our system parameters (NA_*ill*_ =0.24, *α* = 7) the compensation achieved with the NDF closely matches the SLM-based compensation with *σ* = 0.5 (Fig. 1(b-e)).

The detection arm of the microscope is standard for LSFM. Objective O2 (Olympus, UMPLFLN, 0.50/20x, water immersion) images the light-sheet plane onto an sCMOS camera (CAM; Hamamatsu, C13440-20CU, ORCA-Flash4.0) via a tube lens (TL) and appropriate fluorescence filters. The field-of-view (FOV) of the camera is 300 × 300 µm.

Airy light-sheet profiles in the presence of attenuation provided by absorbing NIR dye (American Dye Source, Inc., ADS795WS, absorption coefficient: 1.6 × 10^5^ L mol^−1^ cm^−1^, 2mM) were visualized in fluorescein. Figure 1 (b) shows the Airy light-sheet profile with no attenuation-compensation. Figures 1 (c, d) show the Airy light-sheet with NDF-based and SLM-based (*σ* = 0.5) attenuation-compensation respectively. The peak transverse intensities, normalized to their corresponding values at *x* = 0, as a function of longitudinal coordinate (Fig. 1(e)) clearly shows the intensity decay in the main lobe of the Airy beam without attenuation-compensation. We recover a nearly uniform intensity along the full extent of the light-sheet using either the NDF- or SLM-based compensation techniques.

Absorption and scattering are two key phenomena that impede deeper penetration of incident optical fields into tissue. While absorption and single-scattering yield exponential decay of the incident light intensity described by the Beer-Lambert law, the increasing contribution of multiple scattering with deeper penetration into the sample may yield strong deviations from the expected exponential decay. Prior studies on attenuation-compensation have only considered compensation of an exponential decay. However, it is possible to control the intensity evolution of the beam arbitrarily, and the decay profile may be compensated for with sufficient characterization of the specimen. Using MCmatlab [15], an open-source Monte Carlo radiative transport program, we found that the intensity decay of the incident light field followed an exponential profile even in the presence of high scattering anisotropy (see **Supplementary Note 1**). This result means that compensation of the exponential intensity decay of light is sufficient, and arbitrary control of the intensity evolution of the beam is not required, for a wide range of specimens. Therefore, we are able to use a standard optical element, the NDF, for providing attenuation-compensation of the field.

## 3. TWO-PHOTON AIRY LSFM IMAGING RESULTS

The two-photon Airy LSFM was set up as described in Section 2 to directly compare the image quality achieved between NDF- and SLM-based attenuation-compensation methods. A 3D suspension of 400 nm diameter green fluorescent beads was made in a 1% agarose gel containing 2 mM NIR dye to yield attenuation by absorption. **Supplementary Fig. S2** shows recorded images of these samples. Figure 2 shows line profiles taken through maximum intensity projections of the recorded images, showing an enhancement in signal-to-background ratio (SBR) at depth when attenuation-compensation is used. Both NDF- and SLM-based methods achieved similar enhancements.

**Fig. 2.**
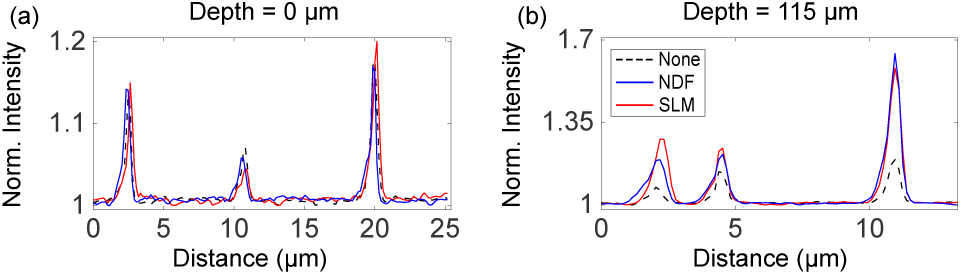
Normalized intensity profiles plotted from recorded two-photon Airy LSFM images of 400 nm diameter fluorescent microspheres in attenuating media (see **Supplementary Fig. S2**). (**a**) Normalized intensity profiles along a line near the start of the FOV where intensities of all three cases match and (**b**)Normalized intensity profiles along a line, 115 µm deeper into the sample.

We further performed a comparison between attenuation-compensation methods in a thick biological specimen of diameter between 300 − 450 µm exhibiting attenuation from both absorption and scattering. Human embryonic kidney 293 (HEK-293) spheroids stably expressing Green Fluorescent Protein (GFP) were imaged. These samples were then fixed in 4% paraformaldehyde and embedded in 1% agarose gel for imaging. Figures 3(a - c) show images acquired with each method and intensity profiles through lines (1) and (2) are shown in Fig. 3(d, e). We performed a SNR analysis of these 3D image stacks in the spatial frequency domain, taking the Fourier transform of each *y* − *z* plane as a function of depth (*x*−axis) into the specimen [16]. We identified spectral bands between *f*_*r*_ = 10 − 50%(2NA/*λ*) and *f*_*r*_ = 80 − 100%(2NA/*λ*) corresponding to “signal” and “noise” respectively, and summed the spectral magnitudes within these bands (see **Supplementary Note 4** for more details). The trend in SNR across the FOV is shown for each method in Fig. 3(f), and the ratio of SNR in NDF- and SLM-based compensation relative to the case of no compensation is shown in Fig 3(g). Beyond a depth of 200 µm, these plots consistently show increases in SNR of ∼33% with NDF- and ∼14% with SLM-based attenuation-compensation, on average, in biological specimen, yielding similar image quality. Besides, a total of 8 HEK-293 spheroid image stacks were acquired, and increases in SNR between 15 − 45% and 5 − 25% in the NDF- and SLM-based attenuation-compensation cases were observed at a depth of 200 µm.

**Fig. 3.**
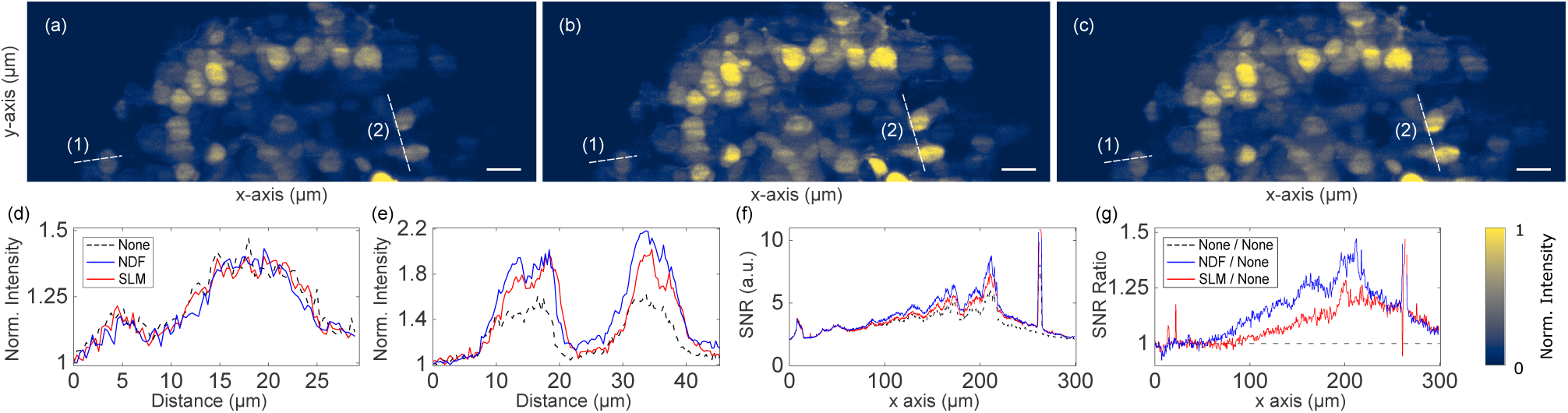
Maximum intensity projections of recorded two-photon Airy LSFM images (a-c) of HEK-293 spheroid stably expressing GFP. (**a**) No attenuation-compensation, (**b**) attenuation-compensation using NDF, and (**c**) attenuation-compensation using SLM (*σ* = 0.5). (**d, e**) Normalized intensity profiles along dashed lines (1) and (2) respectively, shown in (a-c). (**f**) SNR plotted as a function of propagation coordinate, *x*, for no-attenuation compensation, attenuation-compensation using NDF, and attenuation-compensation using SLM (*σ* = 0.5). (**g**) Ratios of SNR with NDF-based compensation (blue) and SLM-based compensation (red) to no compensation. Scalebars: (a-c) 20 µm.

## 4. SINGLE-PHOTON AIRY LSFM IMAGING RESULTS

In addition, we also investigated the performance of attenuation-compensation in the single-photon excitation regime, by substituting the femtosecond laser source for a continuous wave diode laser (Vortran Stradus, *λ* = 488 nm, 150 mW). We used a 1D CPM to generate a 1+1D Airy beam [11, 14] (either a 1D or 2D CPM could be used) and all lenses were replaced with anti-reflection coated versions optimised for 488 nm.

The 1+1D Airy light-sheet profiles in the presence of attenuation were visualized in high concentration fluorescein dye (0.88 mM). Fig. 4 (a-c) show the corresponding Airy profiles with no attenuation-compensaion, NDF-based, and SLM-based (*σ* = 0.8) attenuation-compensation respectively. Uniform intensity in the Airy main caustic after attenuation-compensation can clearly be seen from Fig. 4 (b, c) in contrast to the non-compensated case in Fig. 4 (a). The analogous longitudinal intensity envelopes measured in the above three cases up to an imaging depth of 200 µm are shown in Fig. 4 (d). The difference in area under the curves in this figure quantitatively represents the level of attenuation-compensation provided by both NDF and SLM over their non-compensated counterpart.

**Fig. 4.**
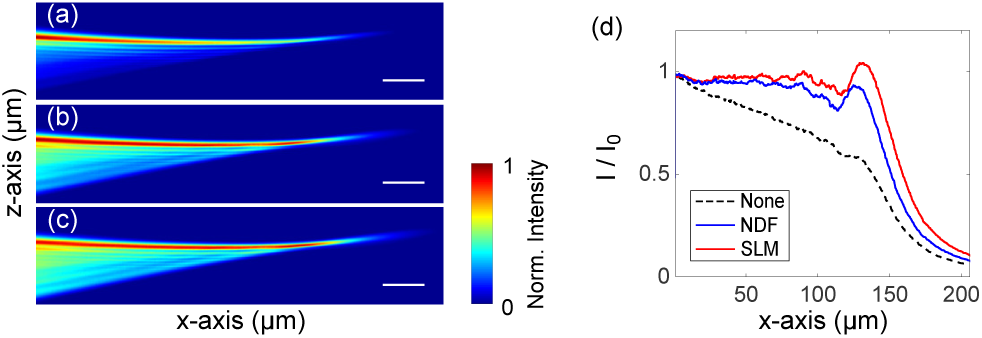
(a - c) show the 1+1D Airy light-sheet profiles visualized in fluorescein (0.88 mM) to show fluorescence and provide attenuation. (**a**) no attenuation-compensation, (**b**) attenuation-compensated by NDF, and (**c**) attenuation-compensated by SLM (*σ* = 0.8). (**d**) Normalized peak transverse intensity versus propagation distance corresponding to beam profiles in (a - c). Scalebars: (a-c) 20 µm.

We made a 3D suspension of 2 µm diameter red fluorescent beads in 1% agarose gel containing 0.88 mM fluorescein, as a phantom acting as an attenuating medium. The attenuation coefficient was determined to be *C*_*attn*_ = 85.9 cm^−1^. **Supplementary Fig. S3** shows deconvolved images of these samples. Figure 5 shows line profiles taken through maximum intensity projections of the deconvolved images [5, 6], showing an enhancement in SBR at depth when attenuation-compensation is used. Both NDF- and SLM-based methods achieved similar enhancements.

**Fig. 5.**
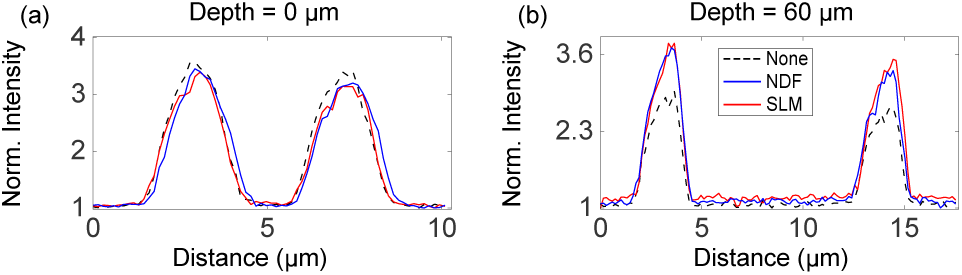
Intensity profiles plotted from deconvolved Airy LSFM images of 2 µm diameter fluorescent microspheres in attenuating media (see **Supplementary Fig. S3**). (**a**) Normalized intensity profiles along a line near the start of the FOV where intensities of all three cases match and (**b**) normalized intensity profiles along a line, 60 µm deeper into the sample.

Finally, we used a biological specimen in the single-photon attenuation-compensated Airy LSFM. Spheroids, comprising of human neuroblastoma (SH-SY5Y) cells stably expressing GFP, were studied. See **Supplementary Fig. S4** for the *x* − *y* maximum intensity projections of the deconvolved images. Similar to the two-photon case, the spheroid was also imaged under three different conditions: no compensation, NDF-based attenuation-compensation, and SLM-based attenuation-compensation(*σ* = 0.8). Similar to the two-photon scenario, SNR corresponding to different compensation schemes was calculated after analysing the Fourier content of the deconvolved image stacks. Fig. 6 (f) plots the SNR against the propagation coordinate at the areas specified in Fig. 6 (a-c) and Fig. 6(g) plots ratios of the data relative to the non-compensated case. On average, these plots show increases in SNR of ∼37% with NDF- and ∼28% with SLM-based attenuation-compensation, in biological specimen, beyond a depth of 120 µm. Additionally, increases in SNR between 25 − 50% and 20 − 40% in the NDF- and SLM-based attenuation-compensation cases were observed at a depth of 140 µm, from a total of 4 SH-SY5Y spheroid image stacks acquired.

**Fig. 6.**
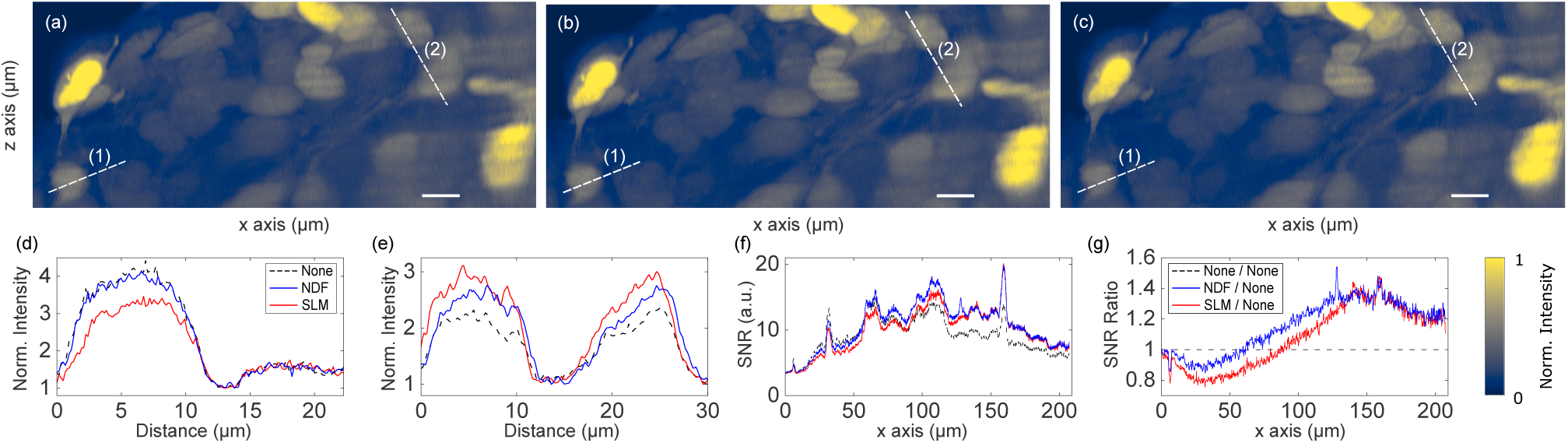
Maximum intensity projections of deconvolved single-photon Airy LSFM images (a-c) of SH-SY5Y spheroid stably expressing GFP. (**a**) No attenuation-compensation, (**b**) attenuation-compensation using NDF, and (**c**) attenuation-compensation using SLM (*σ* = 0.8). (**d**) Normalized intensity profiles along line (1) near the start of the FOV where intensities of all three cases match and (**e**) normalized intensity profiles along line (2), 90 µm deeper into the sample. (**f**) SNR plotted as a function of propagation coordinate, *x*, for no-attenuation compensation, attenuation-compensation using NDF, and attenuation-compensation using SLM (*σ* = 0.8). (**g**) Ratios of SNR with NDF-based and SLM-based compensations to no compensation. Scalebars: (a-c) 10 µm.

## 5. DISCUSSION

We have utilized attenuation-compensation for Airy beam based LFSM to selectively deliver more light deeper into a specimen without increasing the peak illumination intensity, therefore minimizing photo-damage. We have demonstrated the utility of attenuation-compensation provided by a NDF in LSFM and, for the first time, demonstrated an attenuation-compensated two-photon Airy LSFM system. Attenuation-compensation was applied to a light-sheet using an SLM or an NDF, and the performance of the two methods was compared.

Through SNR measurements, we showed an enhanced feature contrast at depth for single- and two-photon Airy LSFM with attenuation-compensation employed using either SLM or NDF. We imaged multiple spheroids of SH-SY5Y and HEK-293 cells with diameter varying between 300 − 450 µm. For brevity, only two sets of data, out of these, are presented here (Figs. 3 and 6). In two-photon Airy LSFM, increases in SNR at 200 µm deep in the HEK-293 spheroid were observed between 5 − 150% and 5 − 50% in the NDF- and SLM-based attenuation-compensation cases respectively. Similar SNR increases of 15 − 65% and 10 − 65% in the NDF- and SLM-based attenuation-compensation cases respectively at 140 µm depth in SH-SY5Y spheroid were observed in single-photon Airy LSFM.

We note that, in the single-photon image data (Fig. 6), attenuation-compensation results in a small reduction in SNR near the start of the FOV which was not observed in the two-photon experiments (Fig. 3). When considering the one- and two-photon light-sheet excitation profiles (Figs. 1 and 4), this can be understood by the considerable background present in the sidelobe region of the beam at the start of the FOV when attenuation-compensation is used. This large background component will reduce the contrast of the sidelobes and have an adverse effect on the deconvolution of the single-photon images. However, in two-photon excitation, the non-linear relationship between illumination intensity and fluorescence excitation will suppress this feature. This interesting effect, coupled with the fact that scattering generally reduces with increasing wavelength and absorption becomes the more dominant loss mechanism at the longer wavelengths used for multi-photon microscopy, suggest that attenuation-compensation may be most useful in the multi-photon regime.

The gold-standard SLM offers dynamic control over the degree of attenuation-compensation. However, in the system we demonstrated, the compensation module in Fig. 1 consisting of an NDF exhibits a virtually identical performance with that of the SLM for the chosen parameters. By altering the optical density range of the linear NDF, the strength of attenuation-compensation can be varied in this configuration. Moreover, combining this with a change in area of illuminated portion on NDF by changing focal lengths of cylindrical lenses leads to a facile pseudo-reconfigurable attenuation-compensation system without the need for complex dynamic diffractive optics.

## ACKNOWLEDGMENTS

We appreciate comments from G Spickermann (M Squared Lasers Ltd) on the manuscript. We thank Dr Wardiya Afshar Saber for providing the cell lines stably expressing GFP.

## DISCLOSURES

The authors declare no conflicts of interest.

See Supplement 1 for supporting content.

