## Supplement 1 for "Multi-photon attenuation-compensated light-sheet fluorescence microscopy"

*Compiled December 12, 2019*

**This document provides supplementary information to "Multi-photon attenuation-compensated light-sheet fluorescence microscopy." This supplementary document consists of 5 supplementary figures and associated notes.**

### SUPPLEMENTARY MATERIALS

#### Supplementary Note 1: Monte Carlo Radiative Transfer Modelling

Absorption and scattering are two major phenomena that impedes deeper penetration of incident optical fields into tissue, but both absorption and single scattering add up to an exponential decay of intensity following the Beer-Lambert law $I/I_0 = \exp(-\mu x)$, where $x$ is the penetration depth and $\mu$ is the tissue attenuation coefficient. The ratio of single scattering to multiple scattering reduces with $x$, and that could have an influence on the attenuation profile of the Airy beam, and subsequently of the light sheet. We employed open-source codebase of MCmatlab [1] for numerically simulating the intensity decay profile of light-field propagating through a standard biological tissue, with varied scattering properties given by the tissue's scattering coefficient $\mu_s$ and its Henyey-Greenstein scattering anisotropy factor $g$. It was observed that the normalized fluence rate exponentially decays against penetration depth over a detection FOV of 300 μm × 300 μm. The MC simulations thus demonstrated that multiple scattering results in negligible deviation from linear attenuation over length scales relevant for microscopy.

Figure S1 presents the results of Monte Carlo simulations. Fig. S1 (a) plots normalized fluence rate per watt of incident light of pencil beam against penetration depth $x$ in tissue with $\mu_s = 50$ and $g$ varying from 0.6 to 1. Fig. S1 (b) plots the same for a tissue with $\mu_s = 100$ and $g$ varying from 0.6 to 1, and Fig. S1 (c) corresponds to a tissue with $\mu_s = 150$ and $g$ varying from 0.6 to 1.

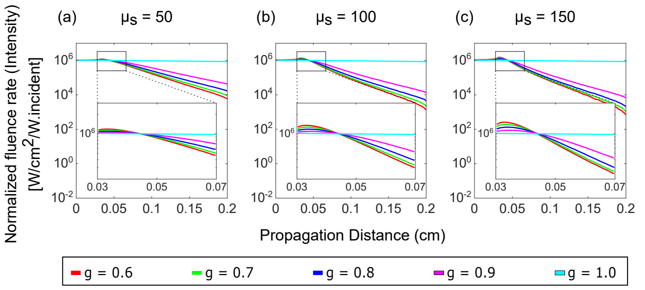


**Fig. S1.** (**a**): Normalized fluence rate per watt of incident light of pencil beam against penetration depth $x$ in tissue with $\mu_s = 50$ and $g$ varying from 0.6 to 1. (**b**): Normalized fluence rate per watt against $x$ in tissue with $\mu_s = 100$ and $g$ varying from 0.6 to 1. (**c**): Normalized fluence rate per watt against $x$ in tissue with $\mu_s = 150$ and $g$ varying from 0.6 to 1. The range of interest 0.03 cm-0.07 cm (400 μm) is highlighted in all the sections.

### Supplementary Note 2: Two-Photon Airy LSM Images

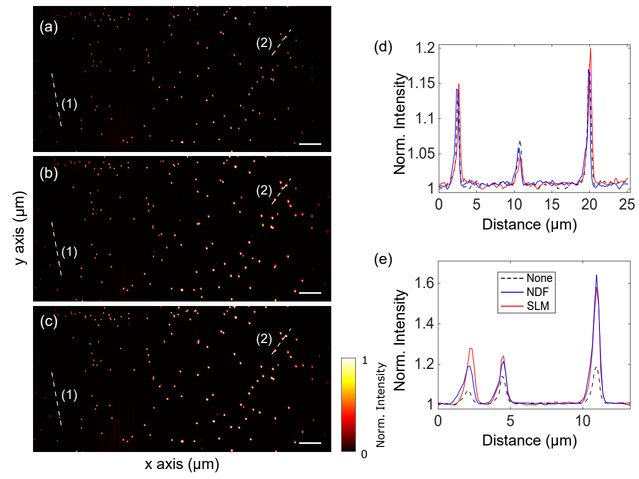


**Fig. S2.** Maximum intensity projections of the recorded two-photon Airy LSFM images (a, b, and c) of 400 $nm$ diameter fluorescent microspheres in attenuating 2 mM concentrated NIR dye solution. (**a**) No attenuation-compensation, (**b**) attenuation-compensation using NDF, and (**c**) attenuation-compensation using SLM ($\sigma = 0.5$). (**d**) Normalized intensity profiles along the dashed line (1) shown in (a), (b), and (c). (**e**) Intensity profiles along the dashed line (2) shown in (a), (b), and (c). Scalebars: (a)-(c) 10 μm.

Fig. S2(a-c) shows the maximum intensity projections of the recorded two-photon Airy LSFM images of 400 $nm$ diameter fluorescent microspheres in 1% agarose gel containing 2 mM attenuating NIR dye, with no attenuation-compensation, NDF-based, and SLM-based ($\sigma = 0.5$) attenuation compensation. The normalized intensity line profiles taken along dashed lines (1) and (2) in Fig. S2(a-c) are reproduced in Fig. S2(d-e) as in manuscript for clarity.

### Supplementary Note 3: Single-Photon Airy LSM Images

Fig. S3 (a-c) shows the maximum intensity projections of the deconvolved single-photon Airy LSFM images of 2 μm diameter fluorescent microspheres in 1% agarose gel containing 0.88 mM concentrated fluorescein dye, with no attenuation-compensation, NDF-based, and SLM-based ($\sigma = 0.8$) attenuation compensation. The normalized line profiles taken along dashed lines (1) and (2) in Fig. S3 (a-c) are reproduced in Fig. S3 (d-e) as in manuscript for clarity.

### Supplementary Note 4: Fourier analysis of Airy LSM images of spheroids

We analysed the 3D image stack of spheroids in the spatial frequency domain and calculated the spectral magnitude within the n$^{th}$ spectral window of the images in y-z plane, $S_n(x)$ [2], given by:

$$S_n(x) = \frac{\int_{f_{r,n-1}}^{f_{r,n}} \int_0^{2\pi} |\tilde{I}(f_r, f_\theta; x)| f_r \, df_r \, df_\theta}{\int_{f_{r,n-1}}^{f_{r,n}} \int_0^{2\pi} f_r \, df_r \, df_\theta} \tag{S1}$$

where $\tilde{I}(f_r, f_\theta; x)$ is the Fourier transform of the image plane $I(y, z; x)$ in cylindrical coordinates and $f_{r,n}$ is the radial spatial frequency separating spectral windows in 10% increments of the diffraction limit: $f_r, n = (n/10)(2NA/\lambda)$. This analysis segments the Fourier space of images orthogonal to depth (y-z) into rings corresponding to different frequency ranges, and calculates the Fourier content in each ring (see Figs. S4 and S5). We observed that the peaks in Fourier spectra corresponding to signal for rings above 40%-50% are strongly attenuated. Besides, since we are looking for nuclei within spheroids with no high frequency details, we could consider the frequency band of 0%-10% as the local dc component. Thus, we considered signal corresponding to 10%-50% Fourier spectra as "signal" and that corresponding to 80%-100% Fourier spectra as "noise", and calculated signal-to-noise ratio (SNR) for the three attenuation-compensation modalities.

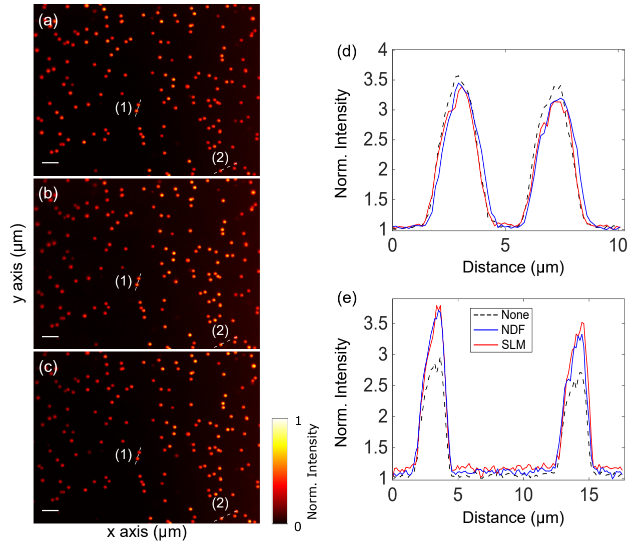


**Fig. S3.** Maximum intensity projections of the deconvolved single-photon Airy LSFM images (a, b, and c) of 2 μm diameter fluorescent microspheres in attenuating 0.88 mM concentrated fluorescein solution. (**a**) No attenuation-compensation, (**b**) attenuation-compensation using NDF, and (**c**) attenuation-compensation using SLM ($\sigma = 0.8$). (**d**) Normalized intensity profiles along the dashed line (1) shown in (a), (b), and (c). (**e**) Intensity profiles along the dashed line (2) shown in (a), (b), and (c). Scalebars: (a)-(c) 10 μm.

### Supplementary Note 5: Spheroids

To generate cellular Spheroids, we used HEK-293 T17 and SH-SY5Y cells stably expressing GFP. A variable number of cells (e.g. 500, 1000, 2000) depending on the wanted spheroids size were plated in ultra-low attachment 96- well round bottom cell culture plates (Corning Costar 7007). After a 48 hours period (or when they have reached the desired size) the spheroids were fixed in PFA. The samples were then embedded in 1% low–melting point agarose and mounted in FEP tubes for the imaging procedure.

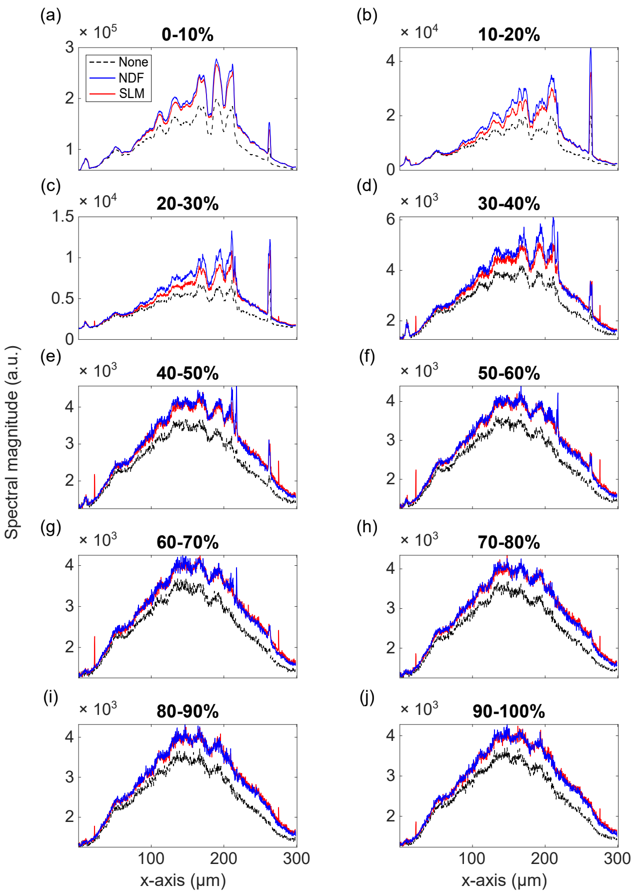


**Fig. S4.** (**a-j**) Plots of spectral magnitude $S_n(x)$ versus tissue depth $x$ for the two-photon Airy LSM recorded image stacks of HEK-293 spheroid stably expressing GFP with no attenuation-compensation, NDF-based, and SLM-based ($\sigma = 0.5$) attenuation compensation.

with airy light-sheet microscopy in cleared and non-cleared neural tissue," Biomed. Opt. Express **7**, 4021–4033 (2016).

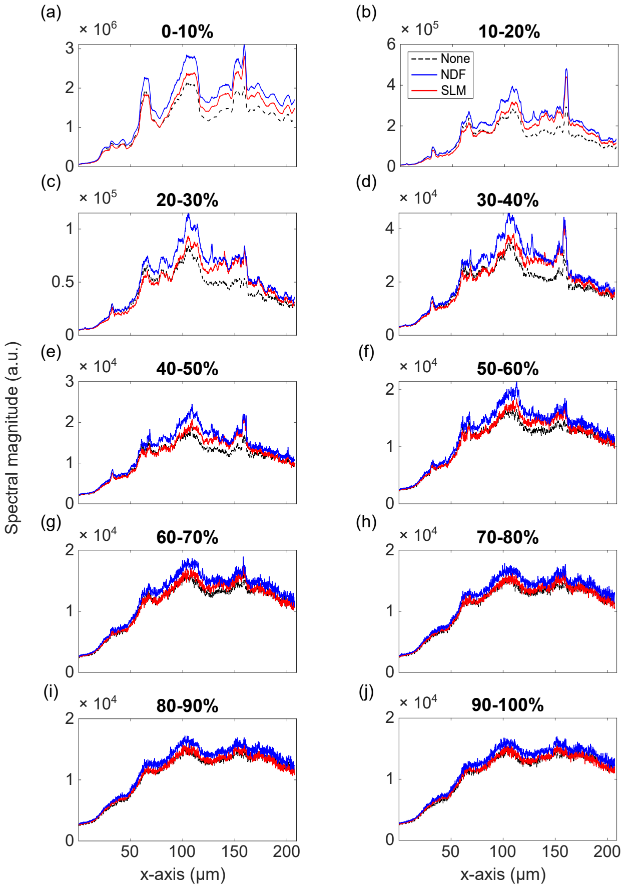


**Fig. S5.** (**a-j**) Plots of spectral magnitude $S_n(x)$ versus tissue depth $x$ for the single-photon Airy LSM deconvolved image stacks of SH-SY5Y spheroid stably expressing GFP with no attenuation-compensation, NDF-based, and SLM-based ($\sigma = 0.8$) attenuation compensation.
